# The synergistic interactions of low-dimensional brain modes

**DOI:** 10.1101/2025.06.19.660516

**Authors:** Ruben Herzog, Jakub Vohryzek, Andrea Luppi, Sebastián Geli, Morten L. Kringelbach, Enzo Tagliazucchi, Gustavo Deco, Yonatan Sanz Perl

## Abstract

Neuroimaging techniques produce vast amounts of data, capturing brain activity in a high-dimensional space. However, brain dynamics are consistently shown to reside in a rather lower-dimensional space, which contains relevant information for cognition and behavior. This dimensionality reduction reflects distinct types of interactions between brain regions, such as redundancy –shared neural information distributed across regions– and synergy, where information emerges only when regions are considered collectively. Significant efforts have been devoted to developing linear and non-linear algorithms to reveal these low-dimensional dynamics, often termed “brain modes.” Here, we apply various dimensionality reduction techniques to resting-state functional magnetic resonance imaging (fMRI) data from 100 healthy participants to examine how synergistic interactions in brain dynamics are preserved by these techniques. We first demonstrate that biologically informed brain parcellation modulates and preserves synergy-dominated interactions. Next, we show that synergy among low-dimensional modes enhances functional-connectivity reconstruction: nonlinear autoencoders not only achieve the lowest reconstruction error but also maximally preserve synergy, outperforming principal component analysis, diffusion maps, and Laplacian eigenmodes. Finally, we confirm previous results suggesting that global signal regression helps to identify synergistic interactions between regions. Our findings establish synergy preservation as a complementary criterion to reconstruction accuracy, highlighting autoencoders as a nonlinear tool for uncovering synergistic low-dimensional brain modes from high-dimensional neuroimaging data.

## Introduction

In the era of big data, dimensionality reduction (DR) techniques have gained increasing attention for their ability to extract meaningful and interpretable information from data. In neuroscience, the high-dimensionality of brain activity recordings is one of the major obstacles hindering experimental and theoretical efforts. Therefore, it is common to transform the data into a reduced set of variables obtained by linear or nonlinear DR methods^1^. In fact, the application of reduction techniques to neural data has a deeper rationale: several works indicate that the true dimensionality of brain activity is smaller than the dimensionality of neuroimaging recordings^2–4^. Indeed, recent results suggest that a set of recurring co-activation motifs obtained from DR of resting-state fMRI reliably predicts individual behavioral traits^5^. Moreover, cognition is enabled by the brain key capacity to dynamically integrate functionally specialized brain regions into a coherent whole^6,7^. Thus, this brain’s integrative property is reflected in the spontaneous self-organization of brain activity into a discrete number of large-scale states or *modes*^8^, which can be associated with specific behavioural and cognitive capacities ^6,9^. This suggests that brain activity involved in cognitive processing is strongly redundant and reduces the effective dimensionality from which emerges the collective behaviour ^10,11^, although its effective scale is still an open question^12^. Indeed, multiple approaches for DR of fMRI data have converged on identifying a small number of “resting state networks (RSN)” in the intrinsic activity of the human brain –whether obtained from Independent Components Analysis (ICA), or clustering of correlations between regions’ temporal fluctuations ^13,14^. These RSNs also appear to have a clear functional role in terms of cognition, as revealed by task-based neuroimaging^15,16^: for example, visual regions are grouped together, and so are motor regions, or regions controlling attention.

Different DR techniques and approaches have been developed over the last few years seeking to compress the high-dimensional signals of brain dynamics^17^. Yet these methods typically optimize variance or local geometry and do not explicitly assess how well they preserve nonlinear and complex interactions across regions. At neuronal level recordings, DR methods have been commonly referred to as “neural manifold learning”^18^, including principal component analysis (PCA), multidimensional scaling (MDS) and Isomap among others. Also, at the level of neuronal circuits, properties of low-dimensional manifolds can represent the state of relevant behavioral variables, such as head direction^19^, and can also be linked to specific circuitry pattern^20^. At the whole-brain level, based on fMRI, different approaches have been proposed to unveil the low-dimensional manifold, resulting, for example, in the so-called brain modes^8^. Laplacian eigenmaps (LEM) –a method that captures the intrinsic geometric structure of the data while preserving local neighborhood relationship^21^– have been used to reveal differences in brain dynamics associated with sleep stages^4^ and with stimuli-induced changes^22^, and also to describe the dynamics across different states of consciousness such as post-comatose patients or altered states^23,24^. In the same direction, autoencoders (AE) –a deep-learning method to compress high-dimensional data into a latent space that preserves data statistical structure– have been used in combination with computational modelling to demonstrate that different states of reduced consciousness can be organized into a one-dimensional manifold embedded into a low-dimensional latent space^25,26^. In sum, DR may not only reveal the effective structure or topology of neural systems^27^, but also provide a more interpretable representation of their activity, from which mechanistic models are built^28^.

DR relies on the assumption that there is redundancy in the data, such that after a given transformation some dimensions can be discarded with only minimal loss of information, enhancing both data compression and interpretation. Despite the usefulness of these techniques, there are continuing discussions on their appropriateness ^29,30^. For instance, it could be the case that variables interact synergistically, which means that there is information that can be obtained if we consider them as a whole functional unit, but not by their isolated examination ^31^. In that case, it is unclear if the transformations performed by DR techniques would leverage and preserve these synergistic information or not. In particular, synergy-dominated interactions in the brain are constrained to mid-to-low orders (i.e. number of interacting regions), while redundancy dominates at higher orders, as expected from the brain large-scale integration^32^. Also, there is evidence suggesting synergistic and redundant information flows within the whole brain^33–36^, which raises the question of to what extent the interactions of the low-dimensional modes leverages and preserves the synergy of the system while avoiding the redundancy of the high dimensional space.

To evaluate whether low-dimensional representations of brain activity preserve these synergistic interactions, and motivated by a recent call for principled compression^5,12^, we leveraged and applied the O-information –a multivariate metric quantifying the balance between synergy and redundancy grounded in information theory^37^. We hypothesize that nonlinear DR methods, particularly AEs, better preserve synergy-dominated interactions among low dimensional modes compared to linear techniques such as PCA or anatomically defined RSN. Additionally, we hypothesize that different parcellations –another way of DR– act as purely spatial coarse-graining and lack any mechanism to preserve nonlinear interdependencies. Consequently, coarser schemes will quench synergy-dominated interactions at lower orders of interaction, whereas finer parcellations will allow synergy to persist to higher orders. This framework’s objective is to offer a principled approach to identify low-dimensional representation that not only minimizes reconstruction error but also retains the inherent synergistic nature of brain dynamics.

## Results

We aimed to investigate the preservation of synergistic interactions in brain dynamics when DR methods are applied to fMRI data. The focus was on comparing the original high-dimensional space of fMRI data with its low-dimensional representations obtained through four different DR techniques (DM, LEM, AE and PCA) and a widely used RSN decomposition^38^ (Figure 1a), as they imply different approaches to reduce brain dimensionality. To achieve this, we employed O-information to quantify the balance between redundancy and synergy (see Methods). A negative O-information implies that the interaction is synergy-dominated, while a positive implies dominance of redundancy. Zero O-information could mean perfect balance or complete independence (i.e. orthogonal variables). Because synergy-dominated interactions tend to be constrained at low-to-mid interaction orders^32^, we computed the O-information across orders of interactions to obtain a more complete characterization of brain function^39^. Analyses were conducted on resting-state fMRI data from 100 participants in the Human Connectome Project (HCP)^40^, using parcellations of varying granularity: Schaefer-200/100^41^, automated anatomical labeling with 90 regions (AAL90)^42^, and Desikan-Killiany with 68 regions (DK68)^43^. Due to the combinatorial explosion that would arise from evaluating all possible high-order interactions for a moderate parcellation with 100 brain regions, we utilized a greedy algorithm^44,45^ to approximate the maximum and minimum O-information across different orders of interaction (Figure 1b, see Methods). In the case of the low-dimensional representation, we used only one parcellation (Schaefer100) as input for all the aforementioned DR techniques using only 7 dimensions to match the number of RSN^38^. Then, all the possible high-order interactions could be computed exhaustively (Figure 1c).

**Figure 1.**
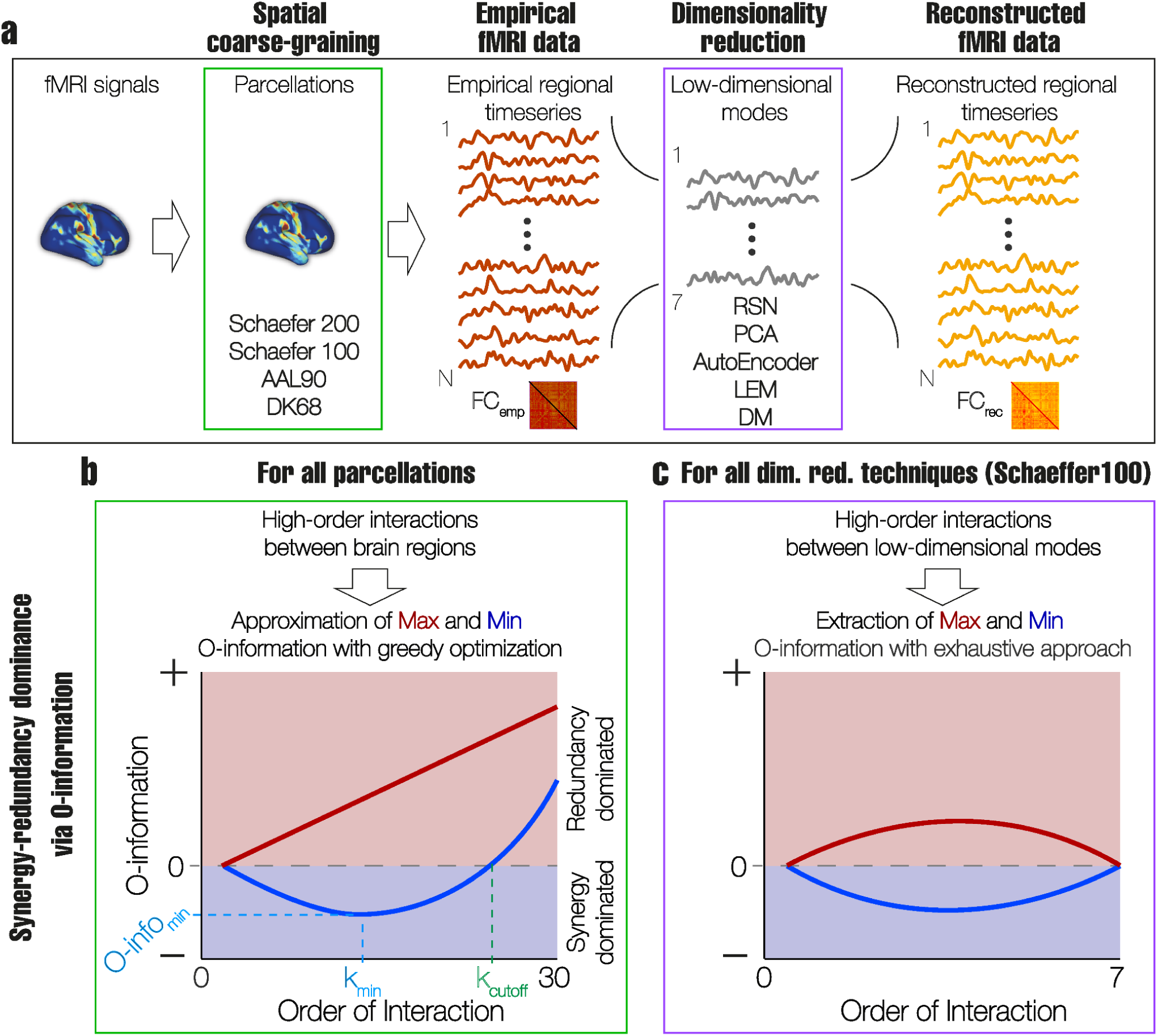
Pipeline overview. **a)** fMRI signals are parcellated (spatial coarse-graining, green square) to produce the empirical signals and their respective functional connectivity matrix (FC_emp_). Then, different DR techniques are used to generate 7 low-dimensional modes to match the number of RSN (purple square). Finally, the fMRI signals and their respective functional connectivity (FC_rec_) are reconstructed from those low-dimensional modes. The reconstruction error is computed as 1-correlation between FC_emp_ and FC_rec_. **b)** The synergy-redundancy balance of high-order interactions is computed via the O-information for all parcellations, considering all orders of interaction, from 3 to 30. To approximate the maximum and minimum O-information across and within orders of interaction, we used a greedy optimization to overcome the combinatorial explosion associated with the high-order approach. The minimum O-information (O-info_min_), its respective order (k_min_) and the order at which synergy no longer dominates interactions (k_cutoff_) are extracted as features to characterize the presence of synergy-dominated interactions on different brain parcellations (depicted in the green square in **a**). **c)** 5 different DR techniques (depicted in the purple square in **a**) are applied to data coarse-grained with the Schaefer100 parcellation. All the possible combinations of high-order interactions between low-dimensional modes are computed from 3 to 7 (exhaustive approach) to assess if synergy is preserved in the low-dimensional space.

### Synergy and redundancy dominance on fMRI resting state data in different parcellations

First, we aimed to characterize the presence of synergy and redundancy in fMRI data parcellated at different granularities, i.e. different number of dimensions (see Methods for details). We found that the maximum O-information increased monotonically with the order of interaction, while the minimum O-information exhibited a convex behaviour (Figure 2a-d), regardless of the parcellation. This convex behaviour for the minimum O-information has been previously shown in fMRI data^32^, and implies that all interactions become dominated by redundancy beyond a certain order. (Figure 2g; at 25.2 ± 4.7 for Schaefer200; 17.3 ± 4.7 for Schaefer100; 18.4±4.8 for AAL90; 14.4±4.8 for DK68). We also found that the average minimum O-information decreased (i.e. more synergy dominance) with the parcellation size (p_FDR_<0.001 for all comparisons Wilcoxon rank sum test after false discovery rate corrected-FDR) (Figure 2e), while the order of interaction at which the minimum O-information was found (i.e. the order at which synergy is more dominant) showed a tendency to decrease with the parcellation size (p_FDR_<0.001 for all comparisons Wilcoxon rank sum test after false discovery rate corrected-FDR)(Figure 2f). Finally, we found that the average ratio between the maximal and minimal O-information tended to remain stable across parcellations (Figure 2h). Consistent with previous results ^32,46^, we showed that fMRI data presented synergy-dominated interactions at small to moderate orders of interactions while redundancy dominated at higher orders.

**Figure 2.**
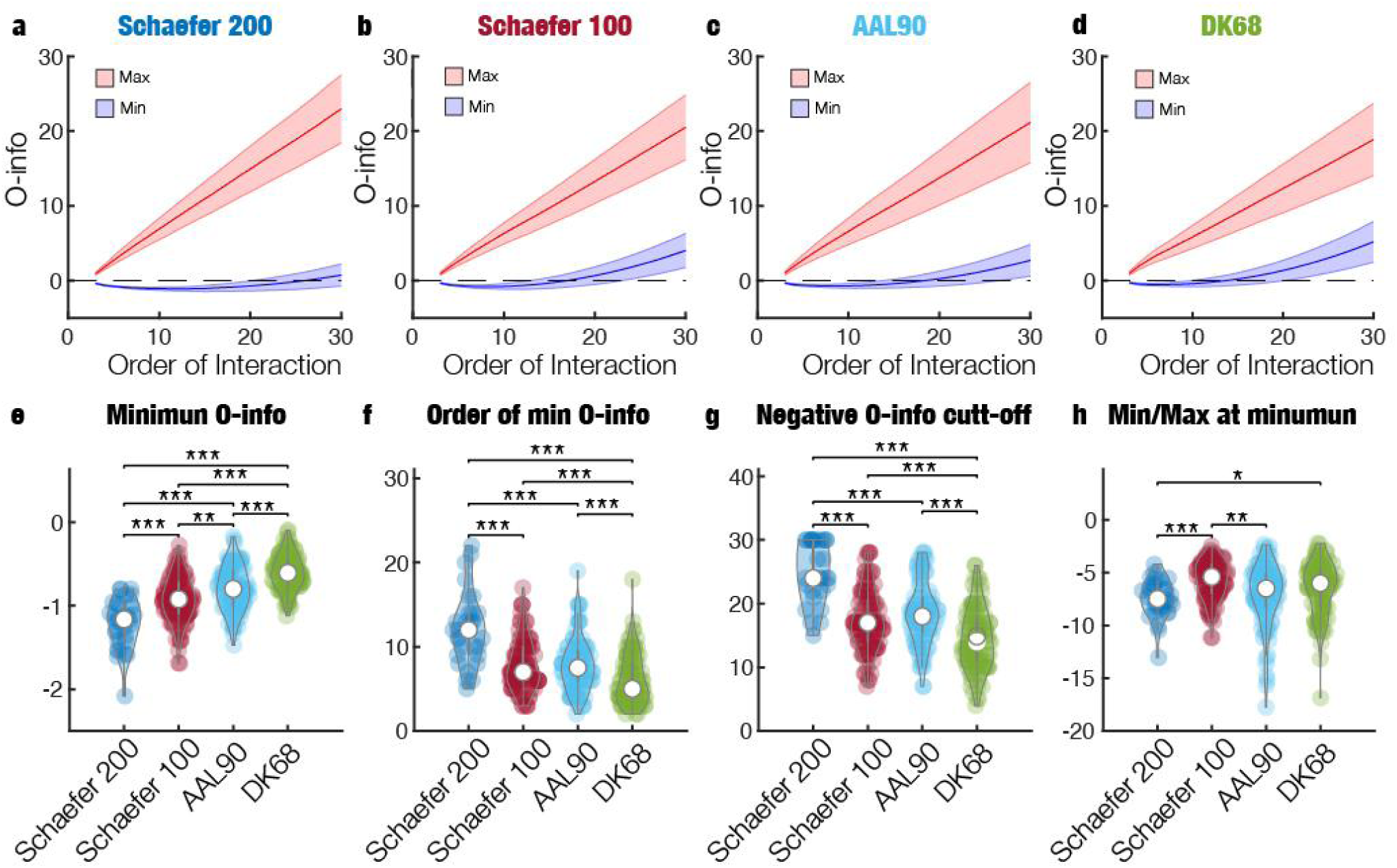
Parcellation size modulates and preserves synergy-dominated interactions in whole-brain fMRI data. **a-d)** Maximum (red) and minimum (blue) O-information at each order of interaction from 3 to 30 for 4 different parcellations (titles). Solid lines are population averages and shaded areas the standard deviation. Note that the minimum O-information exhibits a convex behaviour for all the parcellations. **e)** Violin plots of the minimum O-information for each parcellation. **f)** Violin plots of the order of interaction at which O-information is minimized for each parcellation. **g)** Violin plots of the order of interaction at which O-information is no longer negative for the four parcellations. **h)** Violin plots of the ratio between maximal and minimum O-information at the order where O-information is minimized (violins in **g**).

### Synergy-preserving DR

Synergy refers to information that can be obtained from a system when considering it as a whole functional unit, while not by examining its individual parts alone. Given the prevalence of synergy in brain dynamics^46,33,32^, we argue that methods capable of preserving synergy within low-dimensional modes should retain this key feature of brain dynamics. Furthermore, we hypothesized that synergy-preserving methods would yield comparatively improved reconstructions of brain activity, as more information could be encoded in the same number of dimensions. To test this hypothesis, we evaluated four different DR techniques commonly applied in the analysis of whole-brain fMRI data: PCA, AE, LEM and DM. We focused only on the Schaefer100, as it is a widely used parcellation. For each method, we reduced the data to seven dimensions to facilitate a fair comparison with the widely used Yeo Resting State Network parcellation^38^. We chose to reconstruct the functional connectivity (FC) (Figure 3a), as it is the most widely used feature to represent brain activity, which has been proven useful for characterising different brain states and conditions^47^. Finally, we assessed the reconstruction error by correlating the empirical FC matrix with the one reconstructed from the reduced dimensions (Figure 3b).

**Figure 3.**
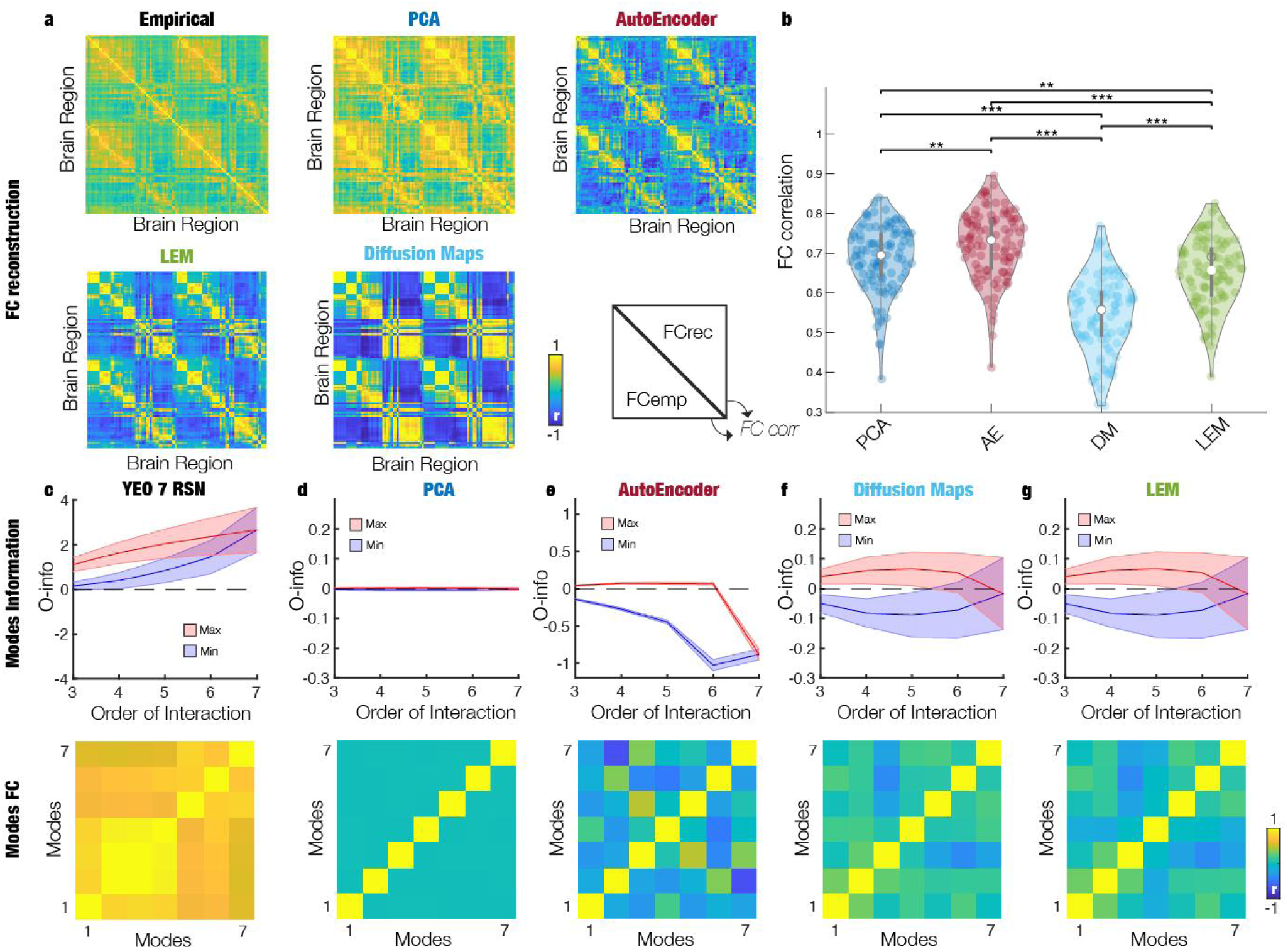
Synergy between low-dimensional modes improves reconstruction of brain functional connectivity. **a)** Functional connectivity matrix for the empirical (Schaefer100 parcellation) and the reconstructed datasets using different DR methods. See Methods for details on the reconstruction. **b)** Reconstruction of the FC matrices, computed as the Pearson correlation between the vectorized upper triangular entries of the empirical and the reconstructed functional connectivity matrix. Each dot represents a subject, and asterisks denote significant differences (** p-val<0.05, *** p-val<0.001). **c-g)** Maximum (red) and minimum (blue) average O-information between the low-dimensional modes obtained for the different methods at each order of interaction. Solid lines represent the average across the population and shaded area its respective standard deviation. Bottom panel shows the pairwise correlations between low-dimensional modes of their respective modes.

Our results showed that the AE provided the best reconstruction accuracy, followed by PCA, LEM, and DM (Figure 3b). Interestingly, this hierarchical organization of methods in terms of reconstruction accuracy remains robust across different dimensionalities of the low-dimensional space (see Supplementary Figure 1).

Then, we analyzed the maximum and minimum O-information at each order of interactions between the low-dimensional modes. PCA, as expected from its orthogonal decomposition approach, produced independent modes with null O-information (Figure 3d). The Yeo 7 RSNs modes, however, were dominated by redundancy, consistent with the high pairwise correlations within resting state networks (Figure 3c). Interestingly, the AE-derived modes exhibited significant synergy dominance at all orders of interactions, even when considering the complete 7-system interaction (Figure 3e). In contrast, DM and LEM produced modes with a balanced contribution of both synergy and redundancy dominated interactions (Figure 3f and 3g), with the overall 7-system interaction yielding an average O-information not significantly different from zero (p-value<0.05), reflecting the minimal pairwise correlations among these modes. These results suggest that while linear decomposition methods like PCA can achieve reasonable reconstructions of whole-brain fMRI data, they fall short in capturing the synergies inherent in brain dynamics due to its orthogonalization procedure (i.e. all PCs are independent). On the other hand, the non-linear decomposition achieved by AEs not only provided the best reconstructions but also effectively preserved the synergistic nature of brain activity. We also investigated the relationship between synergistic and redundant interactions at different orders and the reconstruction accuracy for each method. Importantly, we found no significant correlation across participants for the four methods and all interaction orders after applying FDR correction for multiple comparisons (Supplementary Figure 2). This suggests that both measures provide complementary information about the system.

### Global signal regression enhances the detection of synergistic interactions

Global signal regression (GSR) is a controversial optional step in the preprocessing of fMRI data, to remove global noise and reduce motion-related artifacts, thereby minimizing the influence of widespread, non-neuronal fluctuations on the FC analysis –though possibly at the expense of removing relevant signal from the data^48^. Our hypothesis was that by reducing these common sources of correlation, GSR would decrease redundancy in the data while potentially accentuating synergy-dominated interactions. To test this, we compared our previous results on non-regressed data with those obtained after applying GSR. As expected, we found that GSR reduced the overall values of the FC matrix and enhanced anti-correlations (Figures 4c and 4d). This is as expected from the empirical literature and the mathematics of this operation^49^. We observed a similar convex behaviour in the minimum O-information as in Figure 2 although with a global reduction in both maximum redundancy and maximum synergy (Figures 4a and 4b). Moreover, GSR increased the order of interaction at which the minimum O-information was observed and extended the range of orders where synergy-dominated interactions could be detected (Figures 4e and 4f). These findings suggest that GSR, by reducing shared correlations, diminishes the dominance of redundancy while simultaneously accentuating synergy, as previously observed ^32^. Overall, GSR reduces the dominance of redundancy and enhances the ability to detect synergy-dominated interactions across a broader range of orders of interactions.

**Figure 4.**
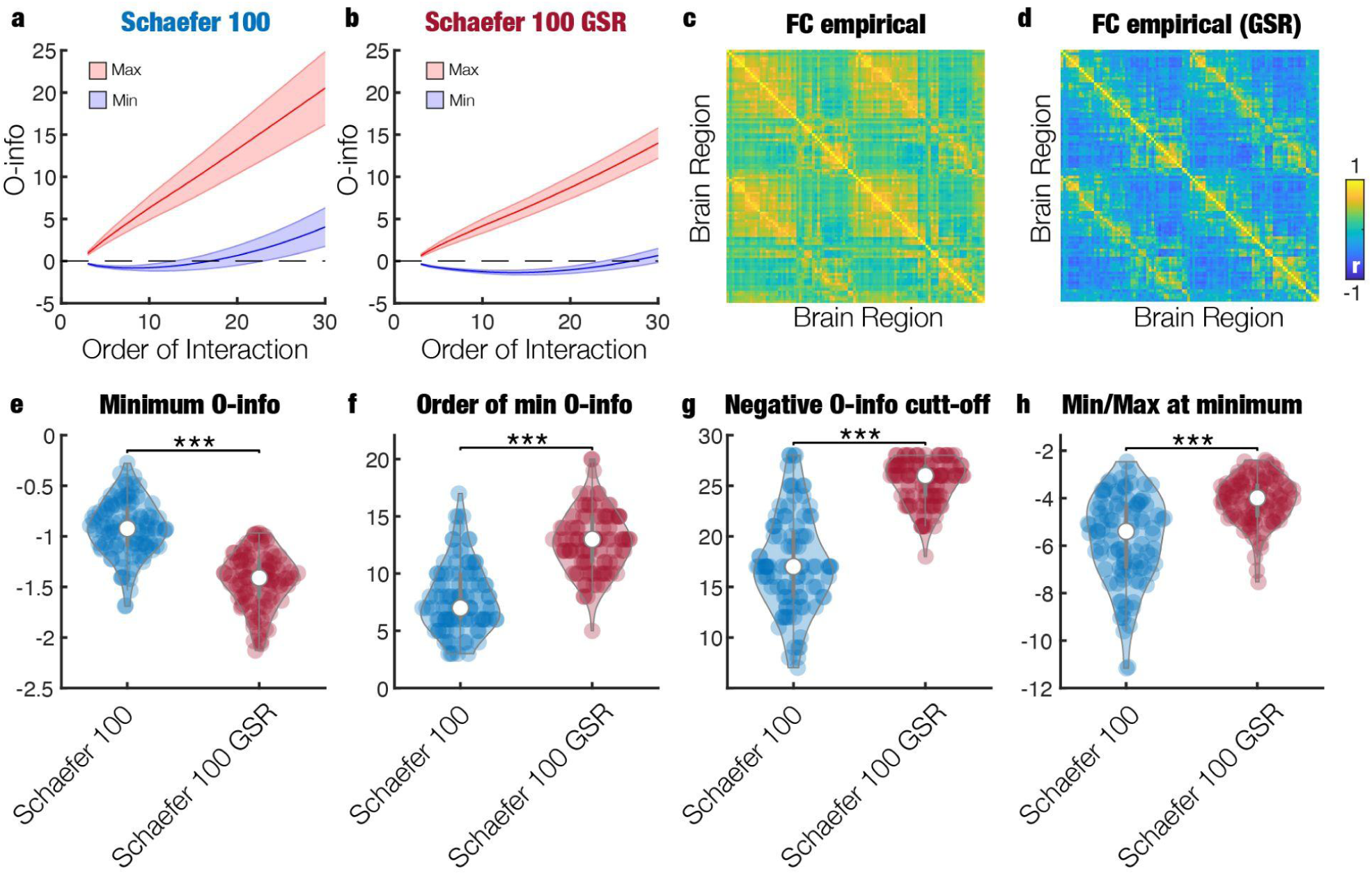
Global signal regression enhances the dominance and range of synergy in whole-brain fMRI data. **a, b)** Maximum (red) and minimum (blue) average O-information for data without GSR and with GSR, respectively. Solid lines represent the average across the population and shaded area its respective standard deviation. GSR induces a global decrease of the O-information values. **c, d)** The respective FC matrices for data without and with GSR, respectively. GSR also induces a global decrease in the FC values, enhancing anti-correlations. **e-f)** Violin plots showing the minimum O-information, its respective order, the order where O-information is no longer negative and the ratio between minimum and maximum O-information at the order shown in **f**, respectively. Overall, GSR decreases the minimum O-information, extends its identifiability across orders of interactions and reduces the max/min O-information ratio. *** p-values<0.001.

Building on the observation that GSR accentuates synergies and reduces redundancy (Figure 4), we investigated whether these changes would influence the performance of DR methods and their ability to preserve synergies between low-dimensional modes. We hypothesized that the qualitative behavior of these methods would remain consistent, while quantitative differences were expected due to the reduction in data redundancy and potential offsetting of values induced by GSR. Consistent with our hypothesis, we observed the same qualitative order of performance across the DR methods, with the AE providing the best reconstruction, followed by PCA, LEM, DM (Figure 5a-b). Regarding maximum and minimum O-information, the qualitative behavior reflected the data without GSR, except for the Yeo 7 Resting State Networks, which exhibited synergy-dominated interactions after GSR (Figure 5c-g). However, the complete 7-variable system was also redundancy-dominated, as for the data without GSR. This result aligns with the amplification of synergies observed in Figure 5 and is consistent with the presence of negative pairwise correlations between the Yeo-derived low-dimensional modes after GSR. For the AE, while the magnitude of both maximum and minimum O-information was reduced, the method still preserved the same qualitative behavior of synergistic interactions, with a dominance of synergy across all orders of interaction, as in data without GSR. Thus, despite the quantitative and qualitative changes that GSR may induce in the FC and the maximum and minimum O-information, AE was still able to generate the best reconstruction while preserving synergies between the low-dimensional modes.

**Figure 5.**
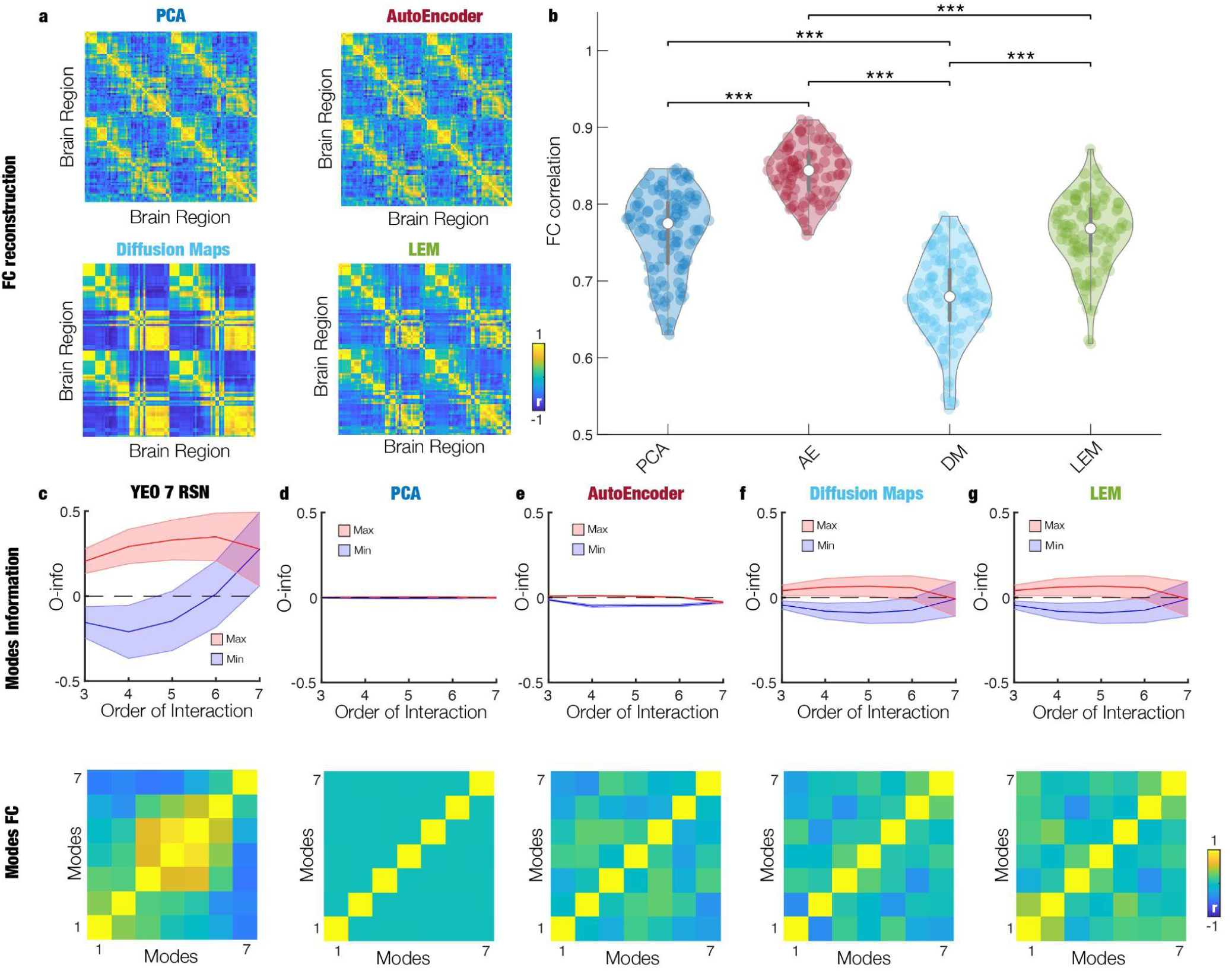
The impact of GSR on the dominance of synergy between low-dimensional modes. **a)** Reconstructed FC matrices with the 4 different DR methods for data with GSR. **b)** Same as in Figure 3b, but for data with GSR. We observe the same qualitative behaviour between methods than for data without GSR. **c-g)** Same as in Figure 3c-g, but for data with GSR. We observe the same qualitative behaviour for all the methods, except for the Yeo approach, where synergy-dominated interactions are observed after GSR. Although for AE the magnitude of the O-information values is reduced, it is still the only method that provides synergy at all orders of interaction. *** p-values<0.001.

## Discussion

This study investigated the capacity of various DR methods to preserve synergistic interactions in whole-brain fMRI data. Synergy, in general terms, refers to information that is in the whole system and that cannot be obtained by analyzing the parts alone. Synergistic interactions have been recently shown to be a key and intrinsic^50^ aspect of brain dynamics^32,33^, tracking aging^46^, changes in consciousness^35,51^ and neurological disorders ^52^. Here we found that synergistic interactions are present in brain dynamics regardless of the parcellation scheme used, confirming previous findings. Moreover, by quantifying synergy and redundancy using O-information^37^, we assessed the trade-offs between preserving synergy and reconstructing FC matrices. Our findings demonstrated that non-linear methods, such as AEs, more effectively preserved both synergy and redundancy, leading to improved FC reconstruction. This work presented a quantitative account for evaluating low-dimensional representations of brain activity not only in terms of reconstruction, but also in terms of preserving synergy, which is lost when using linear methods like PCA.

Our initial approach to DR involved using different brain parcellations, where coarse graining alters the data’s dimensionality based on anatomical and/or functional criteria. This approach offers a biologically plausible reduction compared to purely numerical techniques like PCA or AEs. While the order of synergy-dominated interactions and magnitude of synergy and redundancy dominance were influenced by the parcellation, their ratio remained largely consistent across varying levels of granularity, indicating that these features are intrinsic to brain dynamics. In contrast, non-biologically informed methods (e.g. PCA) that prioritize minimizing redundancy or maximizing independence often sacrifice synergy, leading to an incomplete low-dimensional representation of brain dynamics. To address this, we developed a methodological guide for selecting DR techniques that prioritize synergy preservation, enabling the quantification of retained information in terms of both redundancy and synergy. This framework allows for a more nuanced evaluation of methods beyond reconstruction error by focusing on their capacity to retain key qualitative aspects of brain dynamics.

Regarding the different DR techniques, we start by discussing the RSN method, which is the one with more biological priors. At its origin, RSN analysis was based on independent component analysis (ICA), which aims to maximize the independence between pairs of components^13^. However, we did not perform ICA, rather, we used the well-established clustering-based RSN parcellation provided by Yeo et al (which converged to ICA solutions) ^38^, obtaining results highly dependent on the preprocessing. Without GSR, the RSNs exhibited only redundancy and no synergy. With GSR, synergistic interactions became apparent, suggesting that GSR could help enhance the detection of synergy in brain networks, as previously shown (though possibly at the expense of discarding meaningful biological signals in addition to the noise) ^32^. However, when considering all the reduced dimensions, RSN was redundancy-dominated regardless of GSR, suggesting a sub-optimal use of the low-dimensional space, in comparison with AEs.

PCA (which is related to ICA) is one of the most widely used techniques in different fields, computationally fast, interpretable (e.g. functional gradients ^53,54^), and indeed yields the second-best reconstruction after AE. We speculate that this enhanced performance of PCA relies on its ability to effectively eliminate redundancies, while LEM and DM still exhibit redundancies. However, as recent results suggest, PCA fails at preserving synergy^50^, which we argue is mainly given by diagonalizing the covariance matrix, which produces orthogonal dimensions. The orthogonality constraints have also made PCA insensitive to GSR in terms of the synergy and redundancy between the obtained dimensions.

Both DM and LEM methods are based in spectral graph theory and can account for modular and graded cortical organization^55^. They produced similar results, showing a balance of synergy and redundancy dominated interactions across dimensions. However, they yielded poorer reconstruction errors in comparison with PCA and AEs. Both methods were largely insensitive to GSR, providing consistent decompositions regardless of preprocessing, although they failed at capturing synergy between all the reduced dimensions. Thus, these methods can be suitable for preserving both synergy and redundancy in the reduced space, but with the cost of reduced reconstruction accuracy and more complex computation.

In summary, AEs have outperformed all the methods by preserving both synergy and redundancy and by delivering the best reconstruction of the FC matrix. Even after GSR, the AE maintained a dominance of synergy, though the magnitude of the minimum O-information was reduced. This indicated that AEs were capable of detecting, exploiting and preserving synergy despite changes induced by preprocessing. However, AEs came with drawbacks such as higher computational costs and the need for sufficient data to train the networks. Nonetheless, their ability to preserve synergy across all orders of interaction suggested that preserving synergy might lead to better reconstructions as more information can be encoded within the same number of dimensions.

We also examined how different levels of synergistic and redundant interactions relate to reconstruction performance across methods. Notably, no significant correlations were found across participants for any of the four methods or interaction orders. In particular, the AE is the method that shows more independence between both measures. This indicates that the two types of measures capture distinct and complementary aspects of the system’s behavior.

Our work has several limitations. First, our choice to use seven dimensions was guided by the Yeo 7 Resting State Network parcellation. While this was effective in providing a common comparison framework, as the 7 RSNs are very widely used, it may not be optimal for all datasets, experimental conditions or DR techniques. Future research should investigate more exhaustively the effects of varying the number of dimensions on the preservation of synergy across techniques and experimental conditions. However, by just focusing on resting state data and using the same number of reduced dimensions, we could show significant differences between techniques and provide a guiding principle for further development of DR for brain activity. Regarding measuring the synergy-redundancy balance in higher-dimensional spaces, we employed a greedy optimization approach ^44^. While this method was computationally efficient, it may underestimate the true maximum and minimum O-information. Therefore, the results obtained with the greedy approach should be interpreted as lower and upper bounds, respectively. More computational efficient techniques and optimal usage of high-performance computing architecture are necessary to explore high-order interactions exhaustively, particularly in larger datasets. Despite this limitation, we were able to capture synergistic interactions in a wide range of orders of interactions, regardless of the parcellation. Another limitation of our analysis is that synergistics and redundant interactions were computed without considering time-delayed relationships. We did not consider temporal synergy, which would require methods such as Phi-ID ^56^ to capture time-delayed synergistic interactions. While computing synergy between past and future of the system was out of the scope of this work, we regard including Phi-ID as an interesting avenue for future DR criteria.

Based on the analysis of these different methods, we propose that an optimal DR technique for brain dynamics should fulfill the following requirements: the reduced dimensions i) are synergistic when considered as a whole, maximizing information encoding in the reduced space; ii) exhibit synergy and redundancy across several order of interactions, mimicking the results obtained in the brain’s original space; iii) provide a reconstruction error lower than PCA; and iv) are interpretable in anatomical, functional or cognitive terms. We consider this last requirement an important challenge in whole-brain data analysis, and, to our knowledge, there is no current method that fulfills all these requirements.

Future work should focus on developing autoencoders specifically designed to preserve and exploit the brain synergies. By optimizing this, it may be possible to build models that more accurately reflect the complex interactions underlying brain activity. This would enable improved reconstructions and potentially lead to a deeper understanding of neural processes. A similar criterion could be applied to parcellation techniques, by optimizing them to reduce redundancy while preserving synergy. This could yield a representation of brain activity where regions are maximally independent and, at the same time, they are constrained such that they are still organized as a whole system, following the principle of integration and segregation ^6,7,57^. Such advancements would help to better capture the functional specialization of brain regions while still obtaining a coherent global system. Preserving other features of brain dynamics, such as near-criticality and functional connectivity dynamics, may hold potential for identifying significant changes in brain dynamics^58^. However, their preservation does not inherently enable a synergistic exploitation of the low-dimensional space. Another avenue for future research would be to use these low-dimensional spaces to project brain activity during different states of consciousness, such as sleep, anesthesia, or various neurological disorders. By doing so, it would be possible to investigate how changes in brain dynamics under different conditions are reflected in the synergy and redundancy structure of low-dimensional modes. This could also lead to the identification of biomarkers for specific brain states or conditions and to a potential neurocognitive interpretation of these dimensions. Finally, these DR and synergy-preserving methods should be tested in non-human species. Applying these techniques to animal models could help assess the generalizability of our findings and provide a comparative framework for understanding brain dynamics across species. This could lead to important insights into evolutionary aspects of brain organization and cognitive function.

## Methods

### Neuroimaging data

We used neuroimaging data from using resting-state fMRI recording from *N* = 100 unrelated HCP participants (54 females and 46 males, mean age = 29.1 ± 3.7 years) of the HCP 900 participants’ data release^59^. Data were acquired using as follows: structural MRI: 3D MPRAGE T1-weighted, TR = 2,400 ms, TE = 2.14 ms, TI = 1,000 ms, flip angle = 8°, field of view (FOV) = 224 × 224, voxel size = 0.7 mm isotropic. Two sessions of 15 min resting-state fMRI: gradient-echo EPI, TR = 720 ms, TE = 33.1 ms, flip angle = 52°, FOV = 208 × 180, voxel size = 2 mm isotropic. Here we used functional data from only the first scanning session in left-right (LR) direction.

### Preprocessing

HCP-minimally preprocessed data were used for all participants following Glasser et al. 2013^60^. The minimal preprocessing pipeline is described in detail on the HCP website. The data are preprocessed using the HCP pipeline, which is using standardized methods using FSL (FMRIB Software Library), FreeSurfer, and the Connectome Workbench software^60^. This standard preprocessing included correction for gradient distortions, space and head motion, intensity normalization and bias field removal, registration to the T1 weighted structural image, transformation to the 2-mm Montreal Neurological Institute space, and using the FIX artefact removal procedure^59,61^. The head motion parameters were regressed out and structured artefacts were removed by ICA + FIX processing [independent component analysis followed by FMRIB’s ICA-based X-noiseifier^62^]. Preprocessed time series of all grayordinates are in HCP Connectivity Informatics Technology Initiative (CIFTI) grayordinates standard space and available in the surface-based CIFTI file for each participant for resting state. For the comparison between this minimal preprocessing and the data including global signal regression, we additionally perform a global signal regression to the data.

We used a custom-made MATLAB script using the *ft_read_cifti* function [FieldTrip toolbox^63^] to extract the average time series of all the grayordinates in each region of the different parcellations considering in this work Mindboggle-modified Desikan-Killiany parcellation^43^ with a total of 62 cortical regions (31 regions per hemisphere), Schaefer 100 and 200 (with 50 and 100 cortical regions per hemisphere respectively)^41^ and AAL atlas composed by 90 regions (76 cortical regions and 14 subcortical regions ^42^. which are defined in the HCP CIFTI grayordinates standard space. The BOLD time series were filtered using a second-order Butterworth filter in the range of 0.008 to 0.08 Hz.

### Functional networks

The brain’s functionally interconnected subsystems, where groups of regions consistently activate together, have been a major focus of resting-state functional connectivity research. Intrinsic Functional Networks (IFNs), typically analyzed through correlation techniques, have been repeatedly identified across large-scale resting-state fMRI studies. In this work, we considered the activity of the seven Resting State Networks (RSNs) defined by Yeo and colleagues: Visual, Somatomotor, Dorsal Attention, Ventral Attention, Limbic, Frontoparietal, and Default Mode network^38^. To achieve this, we used the Yeo parcellation mask which is well defined in Schaefer 100 parcellation to associate brain regions to each Yeo 7 RSN. We then average the signals of all regions in each RSN obtaining a time series by network.

### Principal Component Analysis (PCA)

We started using one of the most popular DR techniques, PCA. CA finds the low-dimensional representation Ψ∈ *R*^*Nxp*^, with *p*≪*N* hidden in the original high-dimensional space. This is a linear approach that transforms the data to a low-dimensional space represented by a set of orthogonal components that explain maximal variance. It is fundamentally based on diagonalizing the data covariance matrix. So, the PCA procedure over the timeseries for all brain regions matrix were performed as follows:

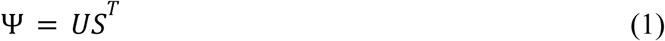

Where U is the left singular vector and S is a diagonal matrix with the singular values of the original input matrix. Importantly, first we compute PCA for the whole group of 100 unrelated HCP participants by concatenating all the time series for each region and participants. We then project the individual time series for each participant to the group’s first *p* = 7 principal components and computed individually the amount of synergy and redundancy interacting in those components.

### AutoEncoder (AE)

Autoencoders are a type of deep neural network commonly used for reducing data dimensionality. These networks are trained to compress input data into a compact representation and then reconstruct the original information with optimal accuracy. The network comprises two main parts: the encoder, which compresses the data into a lower-dimensional latent space, and the decoder, which is a mirror network that reconstructs it back to its original dimensions^64^.

The autoencoder architecture implemented here has eight fully connected layers with dimensions (32, 16, 7, 16, 32), where the latent space dimension, which represents the compressed time series, is 7. Batch normalization and the ReLU activation function were applied to all layers except the middle and output layers. The input data for the network was a 100-dimensional array, representing the state of each node at a given time point. Training was done by minimizing the mean square error (MSE) between the input and output layers using the Adam optimizer. Before training, the data was standardized using z-score normalization and split into 90% training and 10% test sets. The network was trained for 400 epochs or until the validation error plateaued for 50 consecutive epochs. This was implemented in Python using the PyTorch library.

The final result of this training is a low-dimensional representation of the subjects, which was then used to calculate the amount of synergy and redundancy in their interactions.

### Diffusion Maps (DM) and Laplacian Eigenmodes (LE)

The diffusion maps and Laplacian eigenmodes DRs were calculated using the BrainSpace toolbox https://github.com/MICA-MNI/BrainSpace as implemented in MATLAB, with default parameters for density, similarity kernel, and anisotropic diffusion parameter α ^65,66^

To achieve this, we computed the functional connectivity matrix (FC) as the Pearson correlation between every pair of regional fMRI signals for each participant in the 100 unrelated HCP database. Following prior studies, each matrix was z-transformed and thresholded row-wise to retain only the top 10% of values, ensuring 10% density by preserving the strongest connections in each row. Next, we calculated the normalized cosine similarity on the thresholded z-matrix to create a similarity matrix, which reflects the similarity in whole-brain connectivity patterns between regions. While the FC matrix captures how temporally synchronized pairs of regions are, the similarity matrix indicates how similar two regions are in terms of their overall FC patterns. This similarity matrix serves as the input for the diffusion map embedding, or Laplacian eigenmodes algorithm, consistent with previous research on functional gradients^65,66^. Importantly, diffusion map embedding and Laplacian eigenmodes are a nonlinear manifold learning techniques^67^ that exploits the properties of the graph Laplacian, and is therefore related to harmonic mode decomposition, but in this case performed on functional data rather than using the structural connectome, to reveal its contributions to the functional activations.

The mathematical description and deep interpretation about this non-linear DR techniques can be explored in previous work^67,68^.

We computed the low-dimensional modes using both algorithms based on the group average FC and then we aligned the individual modes using Procrustes analysis to the group average modes to improve comparability and correspondence. We then decomposed the fMRI data for each participant in the first *p* = 7 modes.

At each timepoint, corresponding to one TR, the spatial pattern of cortical activity over brain regions at time *t*, denoted as *P_t_*, was decomposed as a linear combination of the set of harmonic modes φ_*k*_, which *k* = 1, …., 7:

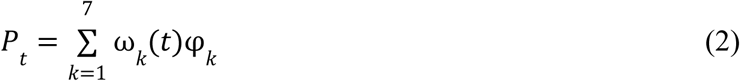

with the contribution ω_*k*_(*t*) of each harmonic mode φ_*k*_ at time *t* being estimated as the projection (dot product) of the fMRI data *P*_*t*_ onto φ_*k*_. We finally considered these projections as the low dimensional representation as in previous works^23,24^ and computed the synergy and redundant interaction between those 7 dimensions for both techniques.

### Computation of O-information

The O-information is a multivariate extension of the Shannon’s mutual information and captures the redundant or synergistic dominant character of a given high-order interaction (3 or more interacting variables). For a multivariate system *X*^*n*^ composed of *n* variables, the O-information (Ω(*X*^*n*^)) is defined as^37^:

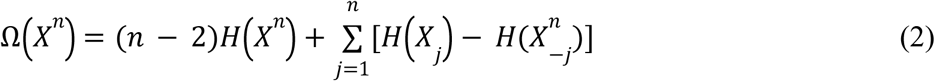

where *H* is the Shannon’s entropy and 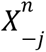 represents the whole system minus the *j-th* variable. When Ω(*X*^*n*^) > 0 the system is redundancy-dominated and when Ω(*X*^*n*^) < 0, the system is synergy-dominated. Estimations of all the entropies were performed using the Gaussian copula approximation ^46,69^. In the case of the DR techniques, all the possible combinations between groups of variables from 3 to 7 were computed and then the maximum and minimum O-information at each order of interaction were extracted. In the case of the original dataset, the number of combinations increases exponentially with the number of nodes, which is known as the combinatorial explosion. For this reason, we used a greedy approach to approximate the maximum and minimum O-information at each order of interaction^45^.

### Greedy algorithm for maximizing and minimizing the O-information

Greedy algorithms perform piece-wise local optimizations to approximate a global optimum. In this case, we wanted to approximate the maximum (minimum) O-information at each order of interaction (from 3 to N, where N is the total number of variables). Previous approaches have used greedy algorithms to find the most affected high-order interactions by neurodegenerative diseases^44^. In detail, we first computed the mutual information for all the possible pairs and used each pair as an initial condition of the search. For a given pair, the algorithm built all the possible triplets that use that pair as a base (so N-2 possible triplets) and selected the one that maximizes (minimizes) the O-information. Then, all the possible N-3 quadruplets that contain that triplet were built and the quadruplet that maximizes (minimizes) the O-information was selected. The algorithm continued like this up to order 30, where no more synergy-dominated interactions were found. To enhance the exploration of the combinatorial space, we took the maximum (minimum) O-information across all the possible pairs used as initial. The algorithm was performed on each subject separately and we reported the population average.

### Functional connectivity reconstruction

The functional connectivity reconstruction using AE is directly obtained from the output of the decoder network. The reconstruction of the original signal using PCA is performed using Eq.1 by pruning the number of components in 7. The reconstructed signal in the Diffusion maps and Laplacian Eigenmodes dimensional reduction can be further defined as follows

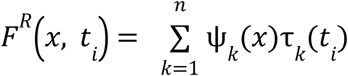

where *F^R^*(*x*,*t_i_*) is the reconstructed signal obtained from spatial components (functional harmonic, FH), ψ_*k*_(*x*), and temporal components (contributions of FH), τ_*k*_ (*t_i_*), at every time point *t_i_*. By projecting the modes onto the time series, it is possible to obtain the contribution of each FH evolving in time. Henceforth, the underlying spatio-temporal activity is described in terms of its spatial mode and temporal dimensions and can be reconstructed back by linear summation.

### Statistical analyses

We applied the Wilcoxon rank-sum method to test the significance and we applied the False Discovery Rate (FDR) at the 0.05 level of significance to correct multiple comparisons^76^.

## Acknowledgments

S.M.G. is an FI fellow with the support of AGAUR, Generalitat de Catalunya and Fondo Social Europeo (2022 FI_B 00511). M.L.K. is supported by the Centre for Eudaimonia and Human Flourishing (funded by the Pettit and Carlsberg Foundations) and Center for Music in the Brain (funded by the Danish National Research Foundation, DNRF117). G.D. is supported by grant PID2022-136216NB-I00 funded by MICIU/AEI/10.13039/501100011033; by ERDF A way of making Europe, ERDF, EU, Project Neurological Mechanisms of Injury, and Sleep-like cellular dynamics (NEMESIS; ref. 101071900), funded by the EU ERC Synergy Horizon Europe; and AGAUR research support grant (2021 SGR 00917) funded by the Department of Research and Universities of the Generalitat of Catalunya. ET is supported by FONDECYT Regular 1220995 (Chile). J.V. and Y.S.P. are supported by the project Neurological Mechanisms of Injury, and Sleep-like cellular dynamics (NEMESIS; ref. 101071900) funded by the EU ERC Synergy Horizon Europe. A.I.L. was supported by St John’s College, Cambridge and the Wellcome Trust (grant number 226924/Z/23/Z). For the purpose of open access, the authors have applied a creative commons attribution (CC BY) licence to any author accepted version arising from this manuscript.

## Author contributions

R.H. and Y.S.P. conceptualized the study, developed the methodology, and drafted the original manuscript; Y.S.P. also performed formal analysis and supervised the project; R.H. and Y.S.P. generated the visualizations; J.V., A.I. L., S.G., M.L.K. and E.T. contributed to writing – review & editing; and G.D. contributed to writing – review & editing and acquired funding.

## Supplementary Material

**Figure Supplementary 1.**
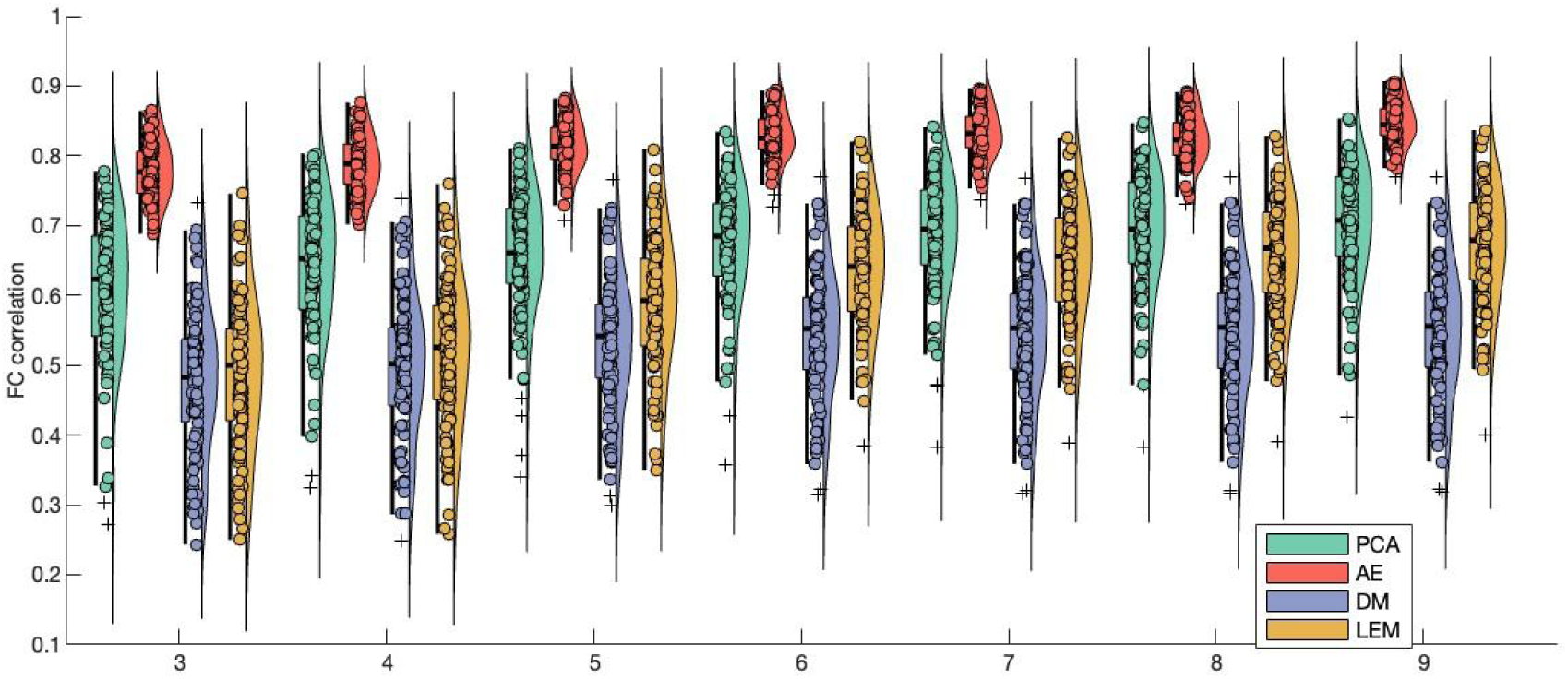
The reconstruction accuracy of the four methods of DR. The reconstruction of the FC matrices using PCA, AE, DM and LEM across the 100 participants as a function of the dimensionality of the reduced space. Importantly, the same hierarchy is observed across dimensions from 3 to 9.

**Figure Supplementary 2.**
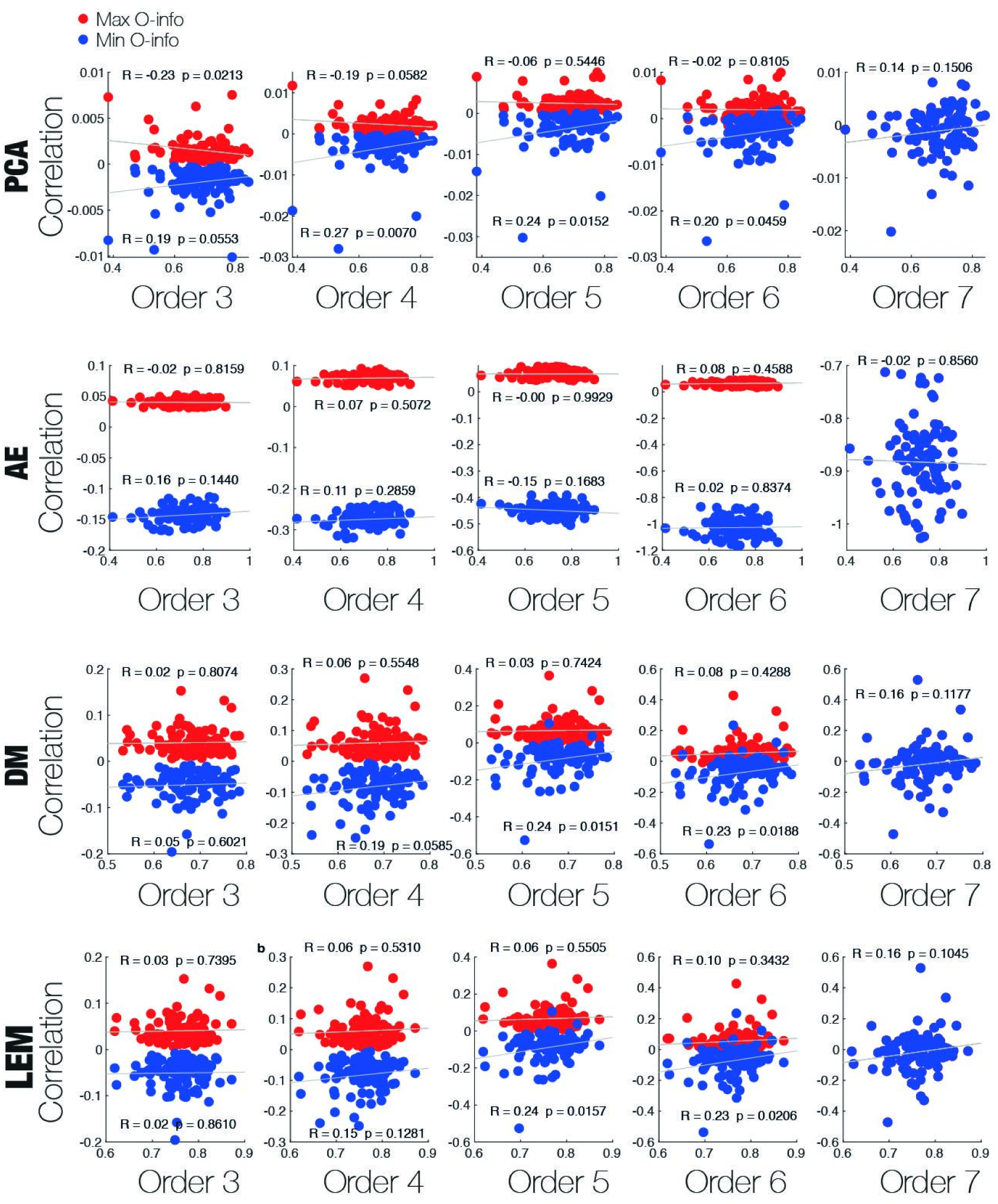
The relation between the minimum and maximum of the O-info and the reconstruction accuracy across orders for the four methods. We investigated the relation between the minimum (blue dots) and maximum (red dot) O-info with the level of FC reconstruction in terms of the correlation across participants. We found no significant correlation for almost all the cases as is indicated in the uncorrected p-values in each scatter plot. However, after FDR multiple correction comparisons within each method no significant correlations are observed. This suggests that both measures are complementary and the level of reconstruction is not determined by the synergistic or redundant interaction at different orders between the low dimensions.

